# Environmental DNA Metabarcoding Effectively Detects Invasive Species, Pests, and Community Changes in Taiwan’s Rice Fields

**DOI:** 10.1101/2025.10.03.680395

**Authors:** Pritam Banerjee, Gobinda Dey, Kathryn A. Stewart, Matthew A. Barnes, Md. Taharia, Mathew Seymour, Chin-Wen Wang, Raju Kumar Sharma, Jyoti Prakash Maity, Chien-Yen Chen

**Affiliations:** Department of Environmental Science, Policy, and Management, University of California, University Avenue and Oxford St, Berkeley, CA, 94720, USA; Department of Earth and Environmental Sciences, National Chung Cheng University, 168 University Road, Min-Hsiung, Chiayi County 62102, Taiwan; Institute of Environmental Sciences, Leiden University, Leiden, 2333 CC, The Netherlands; Department of Natural Resources Management, Texas Tech University, Lubbock, Texas, USA; School of Biological Sciences, The University of Hong Kong SAR, China; Department of Chemistry, School of Applied Sciences, KIIT Deemed to be University, Bhubaneswar, Odisha 751024, India; Center for Nano Bio-Detection, Center for Innovative Research on Aging Society, AIM-HI, National Chung Cheng University, Chiayi 62102, Taiwan

## Abstract

Rice fields represent man-made semi-aquatic wetlands primed for invasive pests. Monitoring rice field biodiversity using conventional methods, however, is time-consuming and laborious. Environmental DNA (eDNA) methods can provide a fast and effective means to monitor rice field communities and inform management decisions. Our study provides proof-of-concept of rice field eDNA biodiversity assessments, with a focus on native and non-native pests across cultivation phases. We collected eDNA samples from locations in southern Taiwan during planting and harvesting, employing eDNA metabarcoding (COI) to detect diverse taxonomic groups. We assigned 77 ASVs across all sites to animal taxa, 34 of which were identified to species. Overall, 18 species were designated as native or non-native (83.3% and 16.6%, respectively), including three major rice pests, *Chilo suppressalis* (native), *Coptotermes formosanus* (native), and *Pomacea canaliculata* (non-native). Cultivation status affected overall diversity, with higher species richness during planting compared to harvesting. No significant differences were observed between native and non-native taxa between cultivation phases. Altogether, we detected a complex environment across trophic levels comprised of both native and non-native agricultural pests using limited sampling effort, demonstrating eDNA analysis as an efficient biomonitoring approach in rice agroecosystems with direct applications for pest, invasive species, and vector surveillance within Taiwan.

## Introduction

Rice fields represent man-made, wetland agroecosystems that support a wide range of biodiversity, from microorganisms to macroinvertebrates and amphibians (Bambaradeniya & Amarasinghe, 2004). Moreover, their alternating hydrologic dynamics, from flooded during the wet seasons to dry conditions in between, mean rice fields house a unique array of biodiversity (Watanabe et al., 2013). Additional ecological characteristics, such as nutrient-rich environments, human influences, and the effects of climate change, collectively make these ephemeral wetlands particularly vulnerable to invasive species (Flanagan et al., 2015). Major threats to the ecological stability and biodiversity in the rice fields include anthropogenic pressures (e.g., intensive monoculture, agrochemical use) and the spread of invasive species (Tilman et al., 2002; Horgan, 2023).

In Taiwan, rice is a major staple crop, deeply embedded in the economy and culture, with almost half of the arable land consisting of rice fields (Hsing, 2014). However, rice fields are one of the most susceptible places for a myriad of invasive pests (Valls et al., 2014). Taiwan’s subtropical climate and extensive irrigation networks further exacerbate invasions by non-native species, posing serious harm to crop productivity and native biodiversity (Barker, 2002; Huang et al., 2010; Wu et al., 2010). Several invasive pest species severely impact rice cultivation in Taiwan; for example, the introduction and establishment of *Pomacea* spp. have become serious threats to rice agriculture (Barker, 2002; Wu et al., 2010; Banerjee et al., 2022), and are currently responsible for large economic losses throughout Asia (Jiang et al., 2022). Migratory rice planthoppers (*Nilaparvata lugens*) (Otuka et al., 2012) and the recently introduced fall armyworm (*Spodoptera frugiperda*) (Tsai et al., 2000) also represent major economic concerns. Compounding impacts, native pests, such as rice stem borers (*Chilo suppressalis*, *Sesamia inferens*, and *Scirpophaga incertulas*), planthoppers (e.g., *Sogatella furcifera*) (Huang et al., 2010), leafhoppers (e.g., *Nephotettix* spp.), and the rice black bug (*Scotinophara coarctata*), additionally contribute to considerable crop damage (Cheng et al., 2010, 2022; Triapitsyn et al., 2021).

Conventional methods of biomonitoring, such as dipnet and funnel traps, are well established and effective for understanding the co-occurrence of organisms associated with rice cultivation (Song & Kuo, 2022), particularly invertebrates. However, conventional methods are also time-consuming and require taxonomic expertise, limiting the scale and scope needed for establishing large-scale temporal monitoring needed to enable adaptive resource management (Baird & Hajibabaei, 2012). Environmental DNA (eDNA)-based metabarcoding has recently revolutionized biodiversity monitoring by enabling the non-destructive detection of targeted or multiple species directly from environmental samples (Taberlet et al., 2012; Valentini et al., 2016; Didaskalou et al., 2022). The adoption of eDNA metabarcoding has proven valuable in agricultural ecosystems, where ecological interactions, including both beneficial and pest organisms, can be monitored for proper management decisions (Kestel et al., 2022; Llanos et al., 2025). However, broad-scale implementation of eDNA-based monitoring in agricultural practices remains limited (Kestel et al., 2022) to a few studies assessing general community observations in rice fields (Ushio et al., 2023; Katayama et al., 2024; Vastano et al., 2024). Previous studies have established species-specific approaches for an early detection method for cryptic invasive gastropods (Banerjee et al., 2022) and fall armyworm (Tsai et al., 2020) in Taiwan. However, more robust and community-level monitoring is necessary for early management action that goes beyond species-specific targets.

Recent advancements in high-throughput sequencing technologies have enhanced the resolution of eDNA metabarcoding approaches, enabling detailed taxonomic and functional assessments of communities present within agricultural environments (Valentini et al., 2016; Seymour et al., 2021; Ushio et al., 2023; Katayama et al., 2024; Vastano et al., 2024). Our study aimed to elucidate a comprehensive understanding of co-occurring species associated with rice field cultivation across two phases of rice cultivation (i.e., planting and harvesting) in Taiwan. To do so, we implemented an eDNA-based metabarcoding approach targeting the mitochondrial cytochrome c oxidase I (COI) gene to investigate the community dynamics of native, non-native, and pest species associated with various rice fields. As environmental conditions (e.g., pH, salinity, temperature, and other abiotic factors) putatively influence the shaping of community composition and function within agroecosystems (Hartmann & Six, 2022) and potentially also modulate eDNA detectability and persistence (Barnes & Turner, 2016; Stewart, 2019), we co-measured water and soil parameters alongside our eDNA sampling.

We hypothesized that (i) eDNA metabarcoding would be an effective method for detecting the vast array of interactive animal species from rice fields via water samples; and (ii) during the planting season, species diversity would be high, but community composition would demonstrate a higher proportion of colonizing non-natives and pests compared to harvesting time due to the drier environment.

## Materials and Methods

### Sample collection and environmental parameter analysis

We collected eDNA samples from eight rice fields in Chiayi County, Taiwan (Figure 1, Table 1). To understand the changes in the community during rice growth, including native and non-native species, all fields were visited two times: (i) planting phase (denoted as samples “PF1-PF8”) which consist of the period of time at the beginning of the growing season where crops are seeded and vegetative growth initiates, and (ii) harvesting phase (“HF1-HF8”), characterized by the ripening phase, with mature rice seedling and dry environment. In our study, planting phase sampling occurred on 22 February 2022, and the harvesting phase sampling occurred on 31 May 2022. We collected five 500 mL surface water samples from each field (N = 40 samples per season). Water samples were kept on ice and transported directly to the Department of Earth and Environmental Sciences, National Chung Cheng University, for analysis.

**Figure 1.**
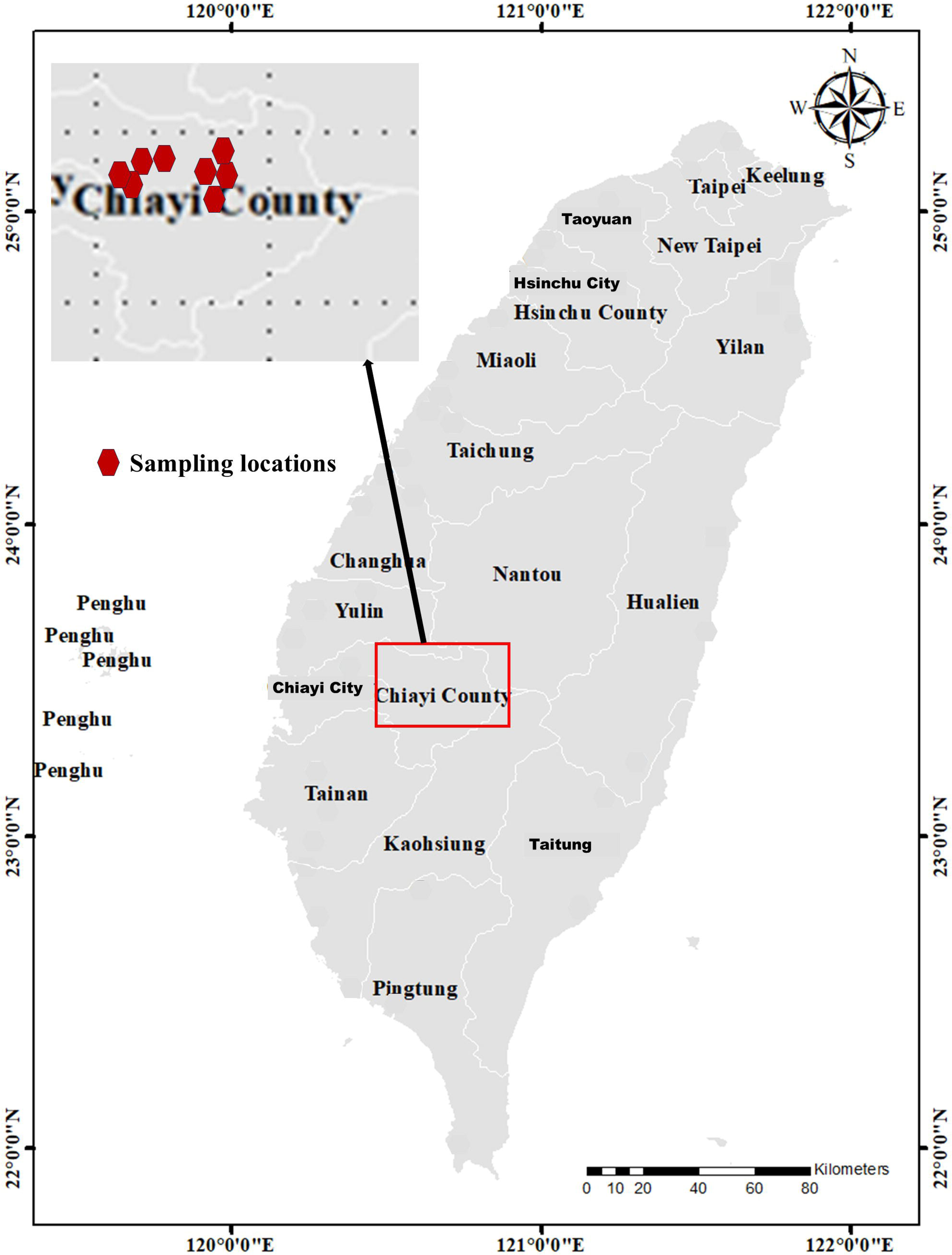
Locations of eight eDNA sampling sites near Chiayi County, Taiwan (February and May 2022)

**Table 1.** Metadata of rice field samples collected at two crop stages. Planting-stage samples (PF1–PF8) were collected on 10 February 2022, and harvesting-stage samples (HF1–HF8) were collected on 31 May 2022, from four rice fields located in Chiayi County, Taiwan. Coordinates for each sampling site are provided.

| Sample name | Field type | Location | Date |
| --- | --- | --- | --- |
| PF1 | planting | 23.567737, 120.454625 | 10-Feb-22 |
| PF2 |  |  |  |
| PF3 |  | 23.540607, 120.380341 |  |
| PF4 |  |  |  |
| PF5 |  | 23.567776, 120.454035 |  |
| PF6 |  |  |  |
| PF7 |  | 23.543891, 120.384877 |  |
| PF8 |  |  |  |
| HF1 | harvesting | 23.567737, 120.454625 | 31-May-22 |
| HF3 |  | 23.540607, 120.380341 |  |
| HF4 |  |  |  |
| HF5 |  | 23.567776, 120.454035 |  |
| HF6 |  |  |  |
| HF7 |  | 23.543891, 120.384877 |  |
| HF8 |  |  |  |

After collection of eDNA, we measured water chemistry at each site including pH, dissolved oxygen (DO) (mg/L), temperature (°C), salinity (PSU), oxidation-reduction potential (ORP) (mV), conductivity (mS/cm), total dissolved solids (TDS) (mg/L), and resistivity (Ω-cm) using a Hanna multiparameter waterproof meter (HANNA instruments, model HI98194, USA), replicating each measurement ten times per site. Subsequently, the water sample was acidified with 5 M HNO to achieve pH < 2, followed by filtration using 0.22-µm nylon syringe filters (13 mm; PureTech™, Taiwan) into 50-ml centrifuge tubes for further heavy metal analysis (see the analytical methods below). Additionally, 3 x 300 g soil samples were collected from each site to measure the physicochemical properties (such as pH and percentage of sand, silt, and clay with texture) and heavy metal concentration, following protocols outlined in Dey et al. (2022). Soil pH was measured using a 1:2.5 soil-to-deionized water suspension with a pH meter. Soil particle size distribution was analyzed by the hydrometer method and classified into textural classes following USDA-NRCS guidelines (Bouyoucos, 1962).

Concentrations of elements Bi, Cd, Mn, Cu, Ni, Co, Cr, Sr, Zn, Pb, and Fe in filtered acid-digested water samples were analyzed using Inductively Coupled Plasma Optical Emission Spectrometry (ICP-OES; Agilent Technologies 5100 ICP-OES, CA, USA). To ensure accuracy and precision, a negative control ultrapure water (PP1-185UV Type I, UNISS, Taiwan) and a positive reference standard were included (ICP multi-element standard solution IV). The recovery rates for the analyzed metals ranged between 94% and 115%, with precision maintained under a 10% relative standard deviation (RSD) for all elements. The soil samples were dried at 70°C for 48 hours, followed by grinding in a mortar and pestle, and then were prepared for heavy metal concentrations analysis using ICP-OES (Agilent Technologies 5100 ICP-OES, USA), based on the method described by Bakshi et al. (2018). Briefly, 0.4 g dried soil was digested with 8 ml aqua regia (a 3:1 mixture of HCl and HNO3) and 2 ml 40% hydrofluoric acid (HF). After cooling, the solution was diluted with 30 mL of deionized water and then filtered with 0.22 µm nylon syringe filters (13 mm; PureTech™, Taiwan) before the ICP-OES analysis.

### eDNA Filtration, Extraction, and Amplification

In the laboratory, all water samples were passed through a GN-6 Metricel membrane mixed cellulose ester filter paper (GN-6 Metricel ®, Pall Corporation, USA) with a diameter of 47 mm and a pore size of 0.45 µm. As rice field water has high TDS, any clogging issues were mitigated by using multiple filter papers as sub-samples (e.g., Hunter et al. 2019). Once our filtration was complete, half of each filter paper was processed for immediate eDNA extraction, and the latter half was preserved at −20 °C in Longmire’s buffer for future use (Longmire et al., 1997). For samples that required multiple sub-filters to mitigate clogging, all filter papers corresponding to the same sample were placed in a single tube for DNA extraction. During the filtration process, three filtration blanks (distilled water) were included to determine contamination and false positive detection during laboratory analysis. Sterilization processes were maintained throughout the workflow, like (i) frequent decontamination with 70% ethanol and 10% bleach, (ii) unidirectional workflow with each step (i.e., filtration, DNA extraction, PCR, and post-PCR) performed in a separate space, and (iii) PCRs were performed in a well-maintained hood.

We extracted eDNA from filters using DNeasy Blood & Tissue kits (Qiagen, Germany) according to the manufacturer’s protocol, but with some modifications as follows: (i) the filter papers were incubated at 60°C overnight in a mixture of 180 µL of ATL lysis buffer and 20 μL Proteinase K; and (ii) the elution step was performed twice, where the spin columns were incubated with the same 50 µL elution buffer at 37°C for 10 minutes (Spens et al., 2017). The quality and quantity of extracted DNA samples were assessed using a Nanodrop spectrophotometer (Implen, Germany) and an Invitrogen Qubit 3.0 fluorometer (Thermo Fisher Scientific, USA). For budgetary reasons, a pooling approach was used for sequencing (Melcher et al., 2024). After extractions, all subsamples (5 samples from each field) were pooled together based on their quantity, thus making 8 pooled samples for each planting (n = 8) and harvesting (n = 8). Finally, these 16 pooled samples were used for further downstream analysis.

A portion of mitochondrial cytochrome c oxidase subunit I (COI) gene was amplified with the following primers: mlCOIintF- 5′- GGWACWGGWTGAACWGTWTAYCCYCC- 3′ and jgHCO2198 - 5′-TAIACYTCIGGRTGICCRAARAAYCA- 3′, specially designed for metazoans (Leray et al., 2013). The reaction mixture of 25 µL for conventional PCR was prepared by mixing 5 µL 5x Fast-Run^TM^ Taq Master Mix with Dye (Protech Technology Enterprise, Taiwan), 0.5 µL each 10 pmol forward and reverse primers (Genomics, Taiwan), 3 µL eDNA sample, and adding high-quality sterile distilled water (Thermo Fisher Scientific, USA) to make up the final volume. For PCR, we tried to add 3 µL eDNA instead of 1µL to allow the potential to amplify all pooled samples. Three PCR replicates for each sample (including blank samples) were carried out. The PCR reactions were performed with the following conditions: an initial denaturation at 94°C for 5 min, followed by a continuous amplification process of 36 cycles, with denaturation at 94°C for 30 sec, annealing at 55°C for 30 sec, and extension at 72°C for 1 min. The final extension was at 72°C for 5 min after the completion of 36 cycles. The PCR products were visualized in a 1.5% agarose gel electrophoresis, and all subsamples/replicates were mixed in equal density ratios (based on band intensity) using the Gel Doc XR system with the QUANTITY ONE*™* software (Bio-Rad, USA). No visible amplification was observed in any blank sample (filtration or extraction blanks); these were therefore excluded from further analysis. The PCR products were then purified using 1.5× SPRI beads (AMPure XP; Beckman Coulter, USA). Purified amplicons were quantified using an Invitrogen Qubit 3.0 fluorometer and submitted for library preparation and sequencing at Genomics Bioscience and Technology Co., Ltd., Taiwan.

### Library preparation and next-generation sequencing

Libraries were prepared using the TruSeq Nano DNA Library Prep Kit (Illumina, USA), and quality was assessed using an Agilent Bioanalyzer 2100 system (Agilent Technologies, USA). The target inserts (∼313 bp of the mitochondrial COI gene fragment) were derived from the PCR amplification described earlier. The final library size, including Illumina adapters, was approximately 450-500 bp, as measured by Bioanalyzer. Libraries were individually indexed (barcoded) during preparation and pooled (multiplexed) in equimolar concentrations before sequencing. Library quantification was performed using qPCR to ensure accurate equimolar pooling. All qualified libraries were sequenced using 300 bp paired-end reads on an Illumina MiSeq platform (Illumina, USA), operated by Genomics Bioscience and Technology Co., Ltd., Taiwan. Sequencing was carried out on a multiplexed standard v3 MiSeq flow cell.

### Data processing

Raw sequencing data were demultiplexed using an in-house script developed by Genomics Bioscience and Technology Co., Ltd., Taiwan, to assign reads to their corresponding samples based on barcode sequences. Trimmomatic (v0.39) was used to remove the adapter sequences, and low-quality bases (QV < 3) using with the following parameters: ILLUMINACLIP:adapter:2:30:10:1:true LEADING:3 TRAILING:3 SLIDINGWINDOW:4:15.

This configuration removes Illumina adapter sequences (allowing up to two seed mismatches), trims low-quality bases from both ends of the reads, and applies sliding window trimming when the average Phred quality score within the window falls below 15 (Bolger et al., 2014). Sequences corresponding to both ends of the mlCOIintF and jgHCO2198 primers were trimmed using Cutadapt (v3.4) (Martin, 2011). Primer trimming was performed with a minimum read length cutoff of 150 bp to remove short, low-information reads, and reads lacking primer sequences were discarded to ensure that only primer-containing sequences were retained. The trimming allowed a minimum overlap of 3 bp and an error rate of 0.1 for primer matching. Denoising, chimera removal, and assignment to Amplicon Sequence Variants (ASVs) were performed using DADA2 (Callahan et al., 2016), following the approach of (Sigsgaard et al., 2021). After error removal, all remaining sequences were then compared against the GenBank nucleotide database (Sayers et al., 2022) (as of September 2022) using BLASTN, and taxa were selected based on the Lowest Common Ancestor (LCA) method (https://github.com/timkahlke/BASTA). The criteria for sequence alignment included a maximum of 500 aligned sequences per query, with a ≥90% query cover and 70-100% sequence similarity. For taxonomic assignments, we applied similarity thresholds as follows: 80-84% for class-level, 85-89% for order-level, 90-94% for family-level, 95-97% for genus-level, and >98% for species-level (Sigsgaard et al., 2021). In cases where multiple taxonomic matches occurred at a given level, the ASV was assigned to the next higher taxonomic rank. The species detected via eDNA metabarcoding were classified as native or non-native based on previous literature and the Catalog of Life in Taiwan (https://taicol.tw/). Furthermore, the ecological roles were documented from previously published literature accessed through Google Scholar, PubMed, and Web of Science.

### Statistical analysis

All statistical analyses were performed using R version 4.3.3 (R Core Team, 2025). Alpha diversity was estimated as observed ASV richness (number of unique ASVs per sample) and visualized across field types using *ggplot2*. Differences in richness between cultivational phases were assessed using Wilcoxon rank-sum tests. Beta-diversity was assessed using non-metric multidimensional scaling (NMDS) based on Bray–Curtis dissimilarities with the metaMDS() function in the vegan package (Oksanen et al., 2001). Group differences in community composition between cultivational stages were tested using PERMANOVA (*adonis2*). To visualize the associations between species and cultivation phases (harvesting and planting), a sankey diagram was constructed using the *networkD3* package (Allaire et al., 2014). All visualization of taxonomic distributions and sample overlaps were performed using the *ggplot2* and *eulerr* packages (Wickham, 2016; Yan, 2021). For the environmental parameter, mean values ± standard error (SE) were calculated, and differences between harvesting and planting periods were evaluated using Welch’s two-sample t-test (α = 0.05).

## Results

### Community-level dynamics of rice fields Sequencing reads and taxonomic composition

After filtering, 1,014,225 sequencing reads were retained, resulting in 67,615 ± 10,984 reads (mean ± standard error; n=15) per sample. Overall, 196 amplicon sequence variants (ASVs) were initially identified across taxa, including Metazoa, Protozoa, Algae, Fungi, and Bacteria. We discarded one sample (HF2) due to low sequencing reads (< 3000). Of the total sequence reads, 54.92% originated from Animalia (80 ASVs), 12.16% from Protozoa (25 ASVs), 4.24% from Bacteria (13 ASVs), 3.29% from Algae (38 ASVs), and 1.40% from Fungi (38 ASVs) (Table 2). Additionally, 23.95% of reads could not be taxonomically assigned with high confidence, as they fell below the 80% similarity threshold (2 ASVs) (Table 2). After removing the unassigned taxa (2 human ASVs were also removed) and targeting animal taxa only (one ASV that could only identify up to Animalia was removed), a total of 77 ASVs were retained for downstream analysis. The majority of reads (93.12%) were from Arthropoda (43 ASVs), followed by Annelida (11 ASVs), Rotifera (8 ASVs), Mollusca (5 ASVs), Chordata (4 ASVs), Gastrotricha (2 ASVs), Cnidaria (2 ASVs), and Nematoda (2 ASVs) (Table 3). In total, 34 species were identified from various groups, and among them, 18 species could be designated as either native or non-native based on published literature, involving 83.3% native, 16.6% non-native (Table 4).

**Table 2.** Taxonomic distribution of amplicon sequence variants (ASVs) across major groups. The table summarizes the total number of reads, ASV richness, and percentage of reads assigned to each group. Unclassified reads with <80% similarity are denoted as “Below 80”.

| Phylum | Total reads | ASVs | Percentage of reads |
| --- | --- | --- | --- |
| Animalia | 557061 | 80 | 54.92 |
| Below 80 | 242931 | 2 | 23.95 |
| Protozoa | 123431 | 25 | 12.16 |
| Bacteria | 43090 | 13 | 4.24 |
| Algae | 33448 | 38 | 3.29 |
| Fungi | 14264 | 38 | 1.40 |

**Table 3.**
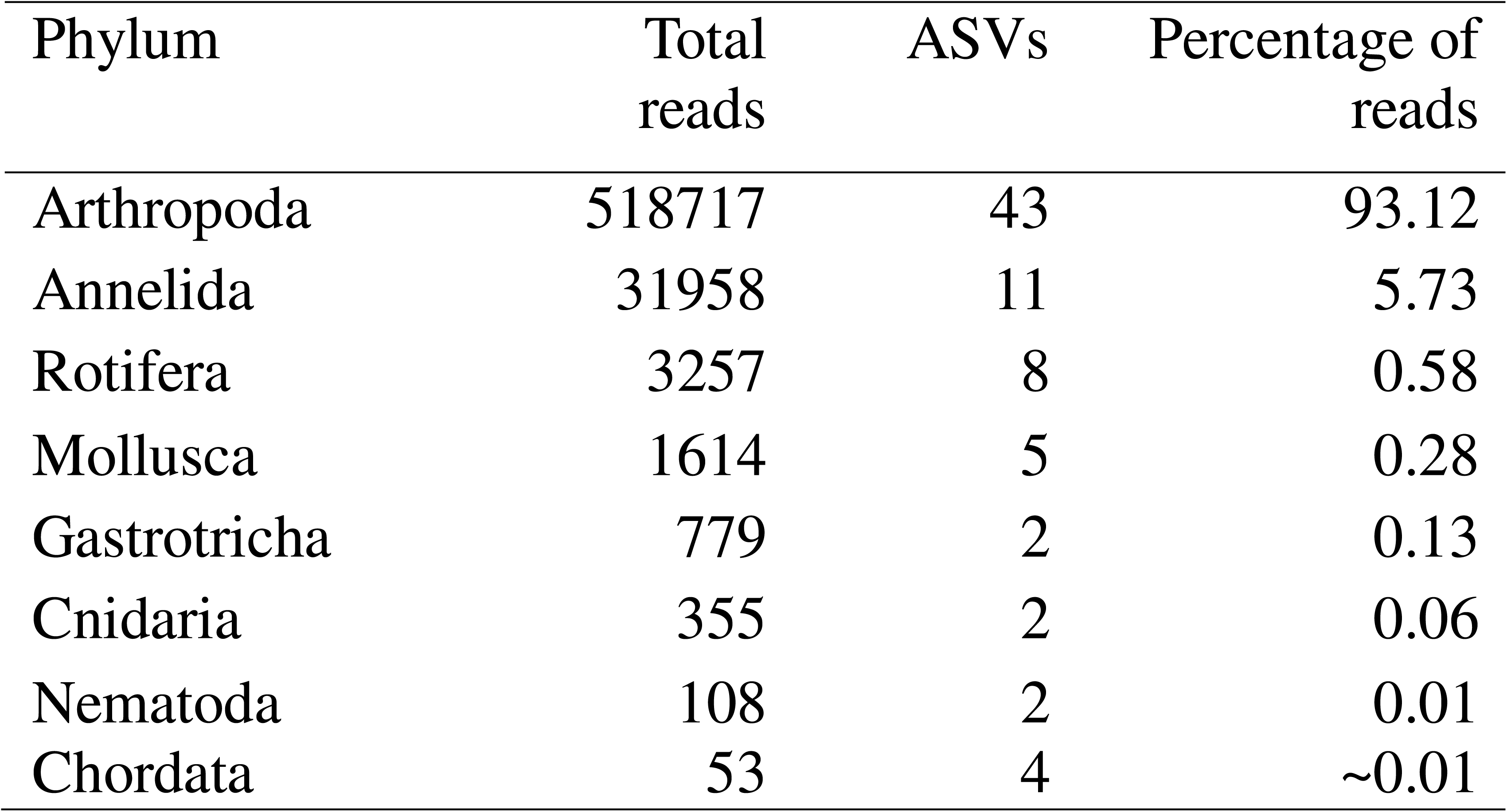
Distribution of Animalia amplicon sequence variants (ASVs) across phyla. The table shows the total number of reads, ASV richness, and relative percentage of reads assigned to each phylum.

| Phylum | Total reads | ASVs | Percentage of reads |
| --- | --- | --- | --- |
| Arthropoda | 518717 | 43 | 93.12 |
| Annelida | 31958 | 11 | 5.73 |
| Rotifera | 3257 | 8 | 0.58 |
| Mollusca | 1614 | 5 | 0.28 |
| Gastrotricha | 779 | 2 | 0.13 |
| Cnidaria | 355 | 2 | 0.06 |
| Nematoda | 108 | 2 | 0.01 |
| Chordata | 53 | 4 | ~0.01 |

**Table 4.**
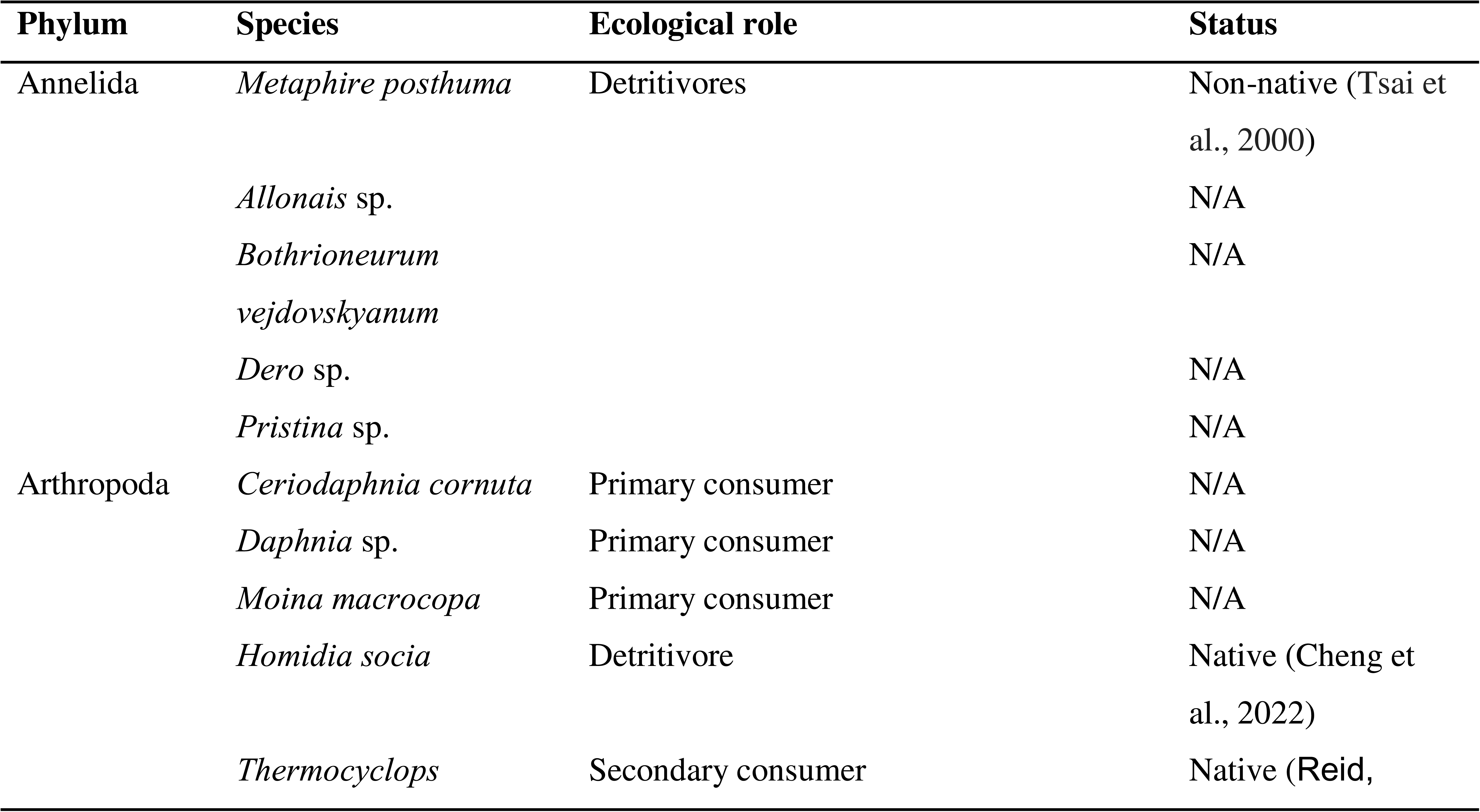
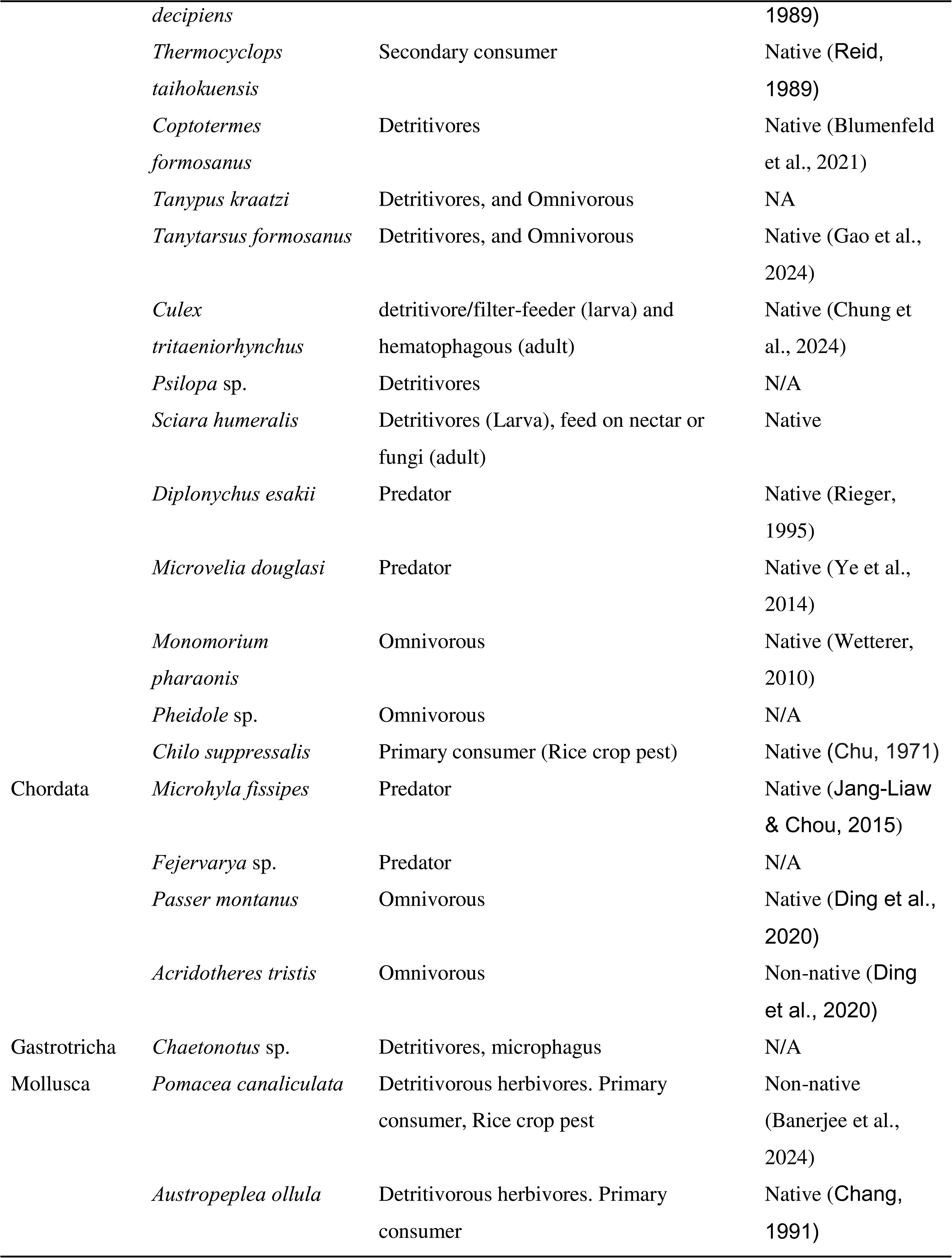
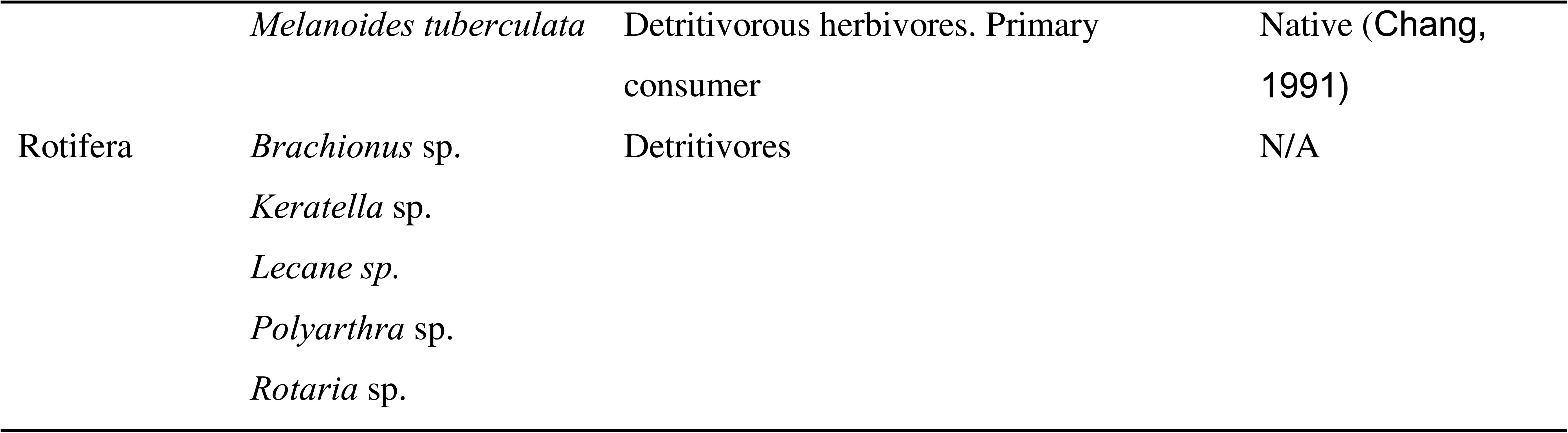
Taxonomic composition, ecological roles, and native/non-native status of organisms detected in Taiwan rice field ecosystems. Species are categorized by phylum, with ecological roles spanning primary consumers, detritivores, predators, and omnivores; status and species type (native, non-native, or not available - N/A) assigned based on references.

### Environmental parameters

Soil pH was slightly alkaline (7.74 ± 0.05 at harvesting, 7.54 ± 0.07 at planting) across sites (Table 5), with textures ranging from sandy loam to loamy sand. Particle size composition remained consistent across both cultivation times, dominated by sand (approximately 72%). Heavy metal concentrations in soil were found to exhibit temporal variation, with Cd, Co, Cr, Cu, Ni, Zn, and Ag generally higher during harvesting, though differences were not statistically significant (Table 5). In contrast, both pH and Lead (Pb) were significantly elevated in soil at harvest (p < 0.05; Table 5).

**Table 5.** The physiochemical properties and heavy metal concentration in agricultural soil during harvesting and planting. p-value are from Welch’s two-sample t-test (α=0.05). a and b subscripts represent significant differences.

| Parameter | Harvesting (Mean ± SE) | Planting (Mean ± SE) | p (Welch t-test) |
| --- | --- | --- | --- |
| pH | 7.74 ± 0.05<sup>a</sup> | 7.54 ± 0.07<sup>b</sup> | **0.0394** |
| Silt (%) | 14.53 ± 1.69<sup>a</sup> | 15.15 ± 1.59<sup>a</sup> | 0.8075 |
| Clay (%) | 13.10 ± 1.65<sup>a</sup> | 12.80 ± 1.47<sup>a</sup> | 0.9016 |
| Sand (%) | 72.37 ± 1.63<sup>a</sup> | 72.05 ± 1.46<sup>a</sup> | 0.8937 |
| Cd (mg/kg) | 0.44 ± 0.07<sup>a</sup> | 0.30 ± 0.06<sup>a</sup> | 0.1694 |
| Co (mg/kg) | 2.41 ± 0.57<sup>a</sup> | 2.02 ± 0.35<sup>a</sup> | 0.6029 |
| Cr (mg/kg) | 70.94 ± 6.48<sup>a</sup> | 64.66 ± 7.12<sup>a</sup> | 0.5536 |
| Cu (mg/kg) | 19.40 ± 1.77<sup>a</sup> | 16.46 ± 2.10<sup>a</sup> | 0.3379 |
| Fe (mg/kg) | 29781.25 ± 2729.60<sup>a</sup> | 24943.06 ± 2848.57<sup>a</sup> | 0.2743 |
| Li (mg/kg) | 37.34 ± 2.49<sup>a</sup> | 37.73 ± 1.77<sup>a</sup> | 0.9074 |
| Mn (mg/kg) | 256.67 ± 32.80<sup>a</sup> | 243.13 ± 36.91<sup>a</sup> | 0.8026 |
| Ni (mg/kg) | 35.11 ± 2.85<sup>a</sup> | 32.98 ± 3.93<sup>a</sup> | 0.6899 |
| Pb (mg/kg) | 35.81 ± 1.39<sup>a</sup> | 29.04 ± 2.49<sup>b</sup> | **0.0492** |
| Sr (mg/kg) | 269.93 ± 34.18<sup>a</sup> | 306.87 ± 27.48a | 0.4489 |
| Zn (mg/kg) | 92.91 ± 9.04<sup>a</sup> | 86.12 ± 7.95<sup>a</sup> | 0.6092 |
| Ag (mg/kg) | 1.26 ± 0.15<sup>a</sup> | 1.14 ± 0.21<sup>a</sup> | 0.6702 |

Water parameters also reflected alkaline pH (7.52 ± 0.05 at harvesting, 7.54 ± 0.05 at planting). ORP was significantly higher during the planting phase (188.99 ± 3.77) compared to the harvesting phase (152.37 ± 11.67; p = 0.0270). Dissolved oxygen (DO) was also higher during planting (5.27 ± 0.26 mg/L) than harvesting (4.45 ± 0.02 mg/L; p = 0.0197). Similarly, conductivity increased during planting (0.55 ± 0.05 mS/cm) relative to harvesting (0.40 ± 0.04 mS/cm; p = 0.0363), and total dissolved solids (TDS) were greater in planting (259.80 ± 22.12 mg/L) than harvesting (187.08 ± 20.39 mg/L; p = 0.0424). In contrast, temperature was significantly higher during the harvesting phase (29.32 ± 0.88 °C) compared to the planting phase (26.08 ± 0.76 °C; p = 0.0231). Among trace metals, only lithium (Li) showed a statistically significant difference, with slightly higher values during harvesting (0.27 ± 0.00 mg/L) than planting (0.26 ± 0.00 mg/L; p = 0.0082). Other parameters, including pH, mV pH, resistivity, PSU, atmospheric pressure, Fe, Mn, and Sr, did not differ significantly between phases (p > 0.05).

### Planting vs harvesting phase

The relative abundance of the top 10 taxa at different taxonomic levels (order, family, genus, and species) varied markedly between harvesting and planting phases (Figure 2). At the order level, Diplostraca was the dominant order in both field types, accompanied primarily by Diptera in harvesting fields and by Crassiclitellata and Cyclopoida in planting fields. At the family level, Moinidae dominated both harvesting and planting fields, co-occurring with Culicidae and Daphniidae in harvesting fields and with Megascolecidae and Cyclopidae in planting fields. At the genus level, *Moina* sp., dominated both harvesting and planting fields, co-occurring with *Culex* sp., and *Ceriodaphnia* sp., in harvesting fields and with *Metaphire* sp., and *Thermocyclops* sp., in planting fields. At the species level, *Moina macrocopa* was dominant in both fields, accompanied by *Culex tritaeniorhynchus* and *Ceriodaphnia cornuta* in harvesting fields and by *Metaphire posthuma* in planting fields.

**Figure 2.**
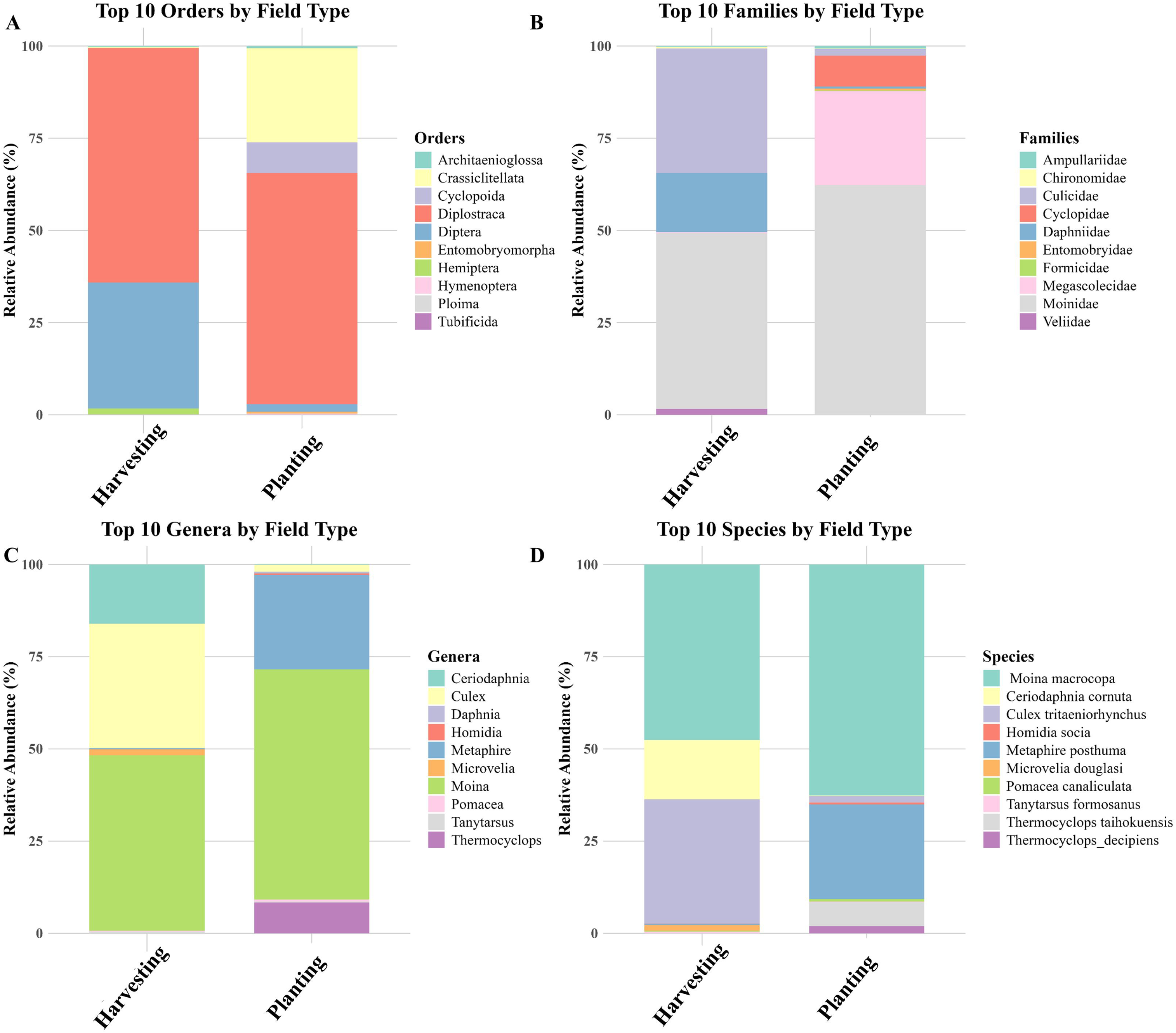
Relative abundance of the top metazoan 10 taxa by field type. Stacked bar plots showing the relative abundance (%) of the top 10 taxa detected in *harvesting* and *planting* field types, displayed at four taxonomic levels: (A) orders, (B) families, (C) genera, and (D) species.

Of note, species overlap between planting and harvesting stages was substantial (44.1%; Figure 3A) but also demonstrated some exclusive taxa between both cultivation phases, with more unique species during the planting (35.7%) than the harvesting phase (14.3%) (Figures 3A, B). This was corroborated by a Sankey diagram showing the association between the cultivation phases and the relative abundance (based on sequencing reads) of taxa (Figure 3B). Moreover, species richness was higher during the planting phase, with 12 native, 3 non-native, and 14 undetermined (status) taxa, compared to 9 native, 2 non-native, and 9 undetermined (status) taxa in the harvesting phase (Figure 4; Table 4). Still, alpha-diversity did not differ significantly between harvesting and planting phases (p-value = 0.52) (Figure 5A). The median species richness was slightly higher during harvesting compared to planting phases, although the variation was larger during planting, with a few high diversity outliers (Figure 5A).

**Figure 3.**
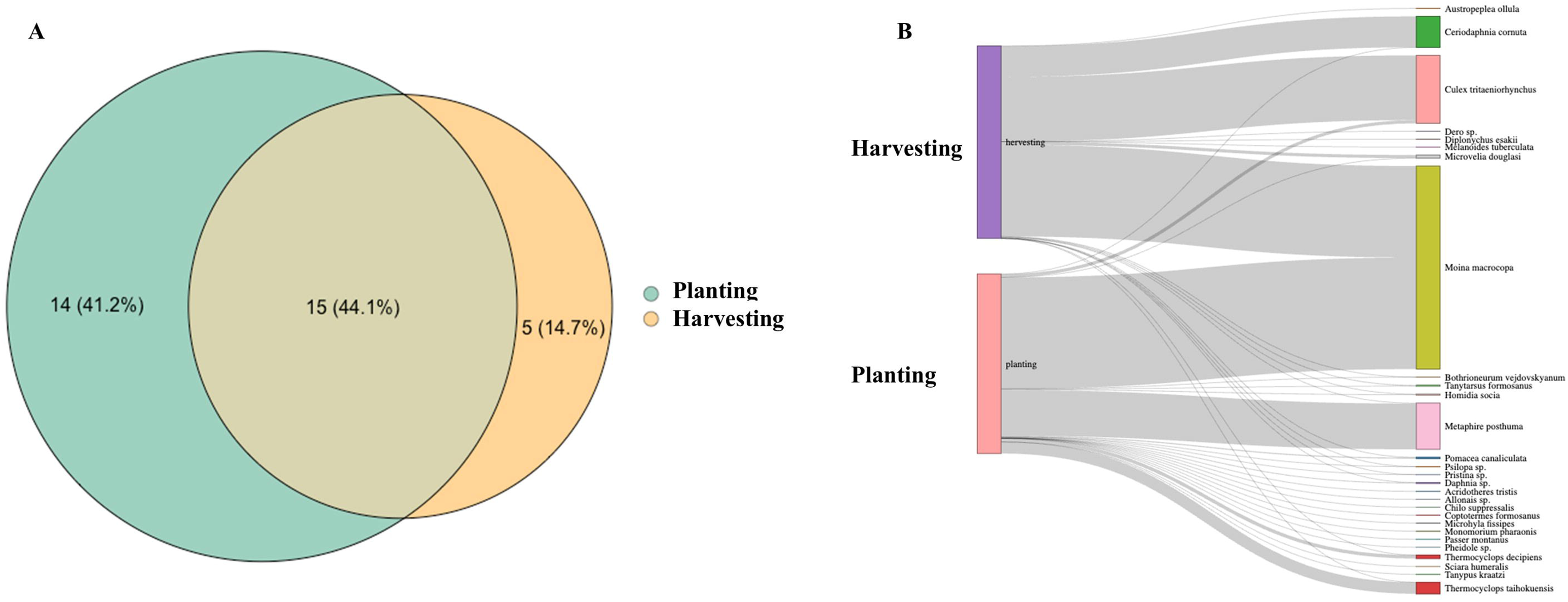
(A) Venn diagram of species detected in planting and harvesting fields. Circle and overlap size denote proportional species diversity representation across field types. (B) Sankey diagram showing the association between rice field types (planting and harvesting) and detected species.

**Figure 4.**
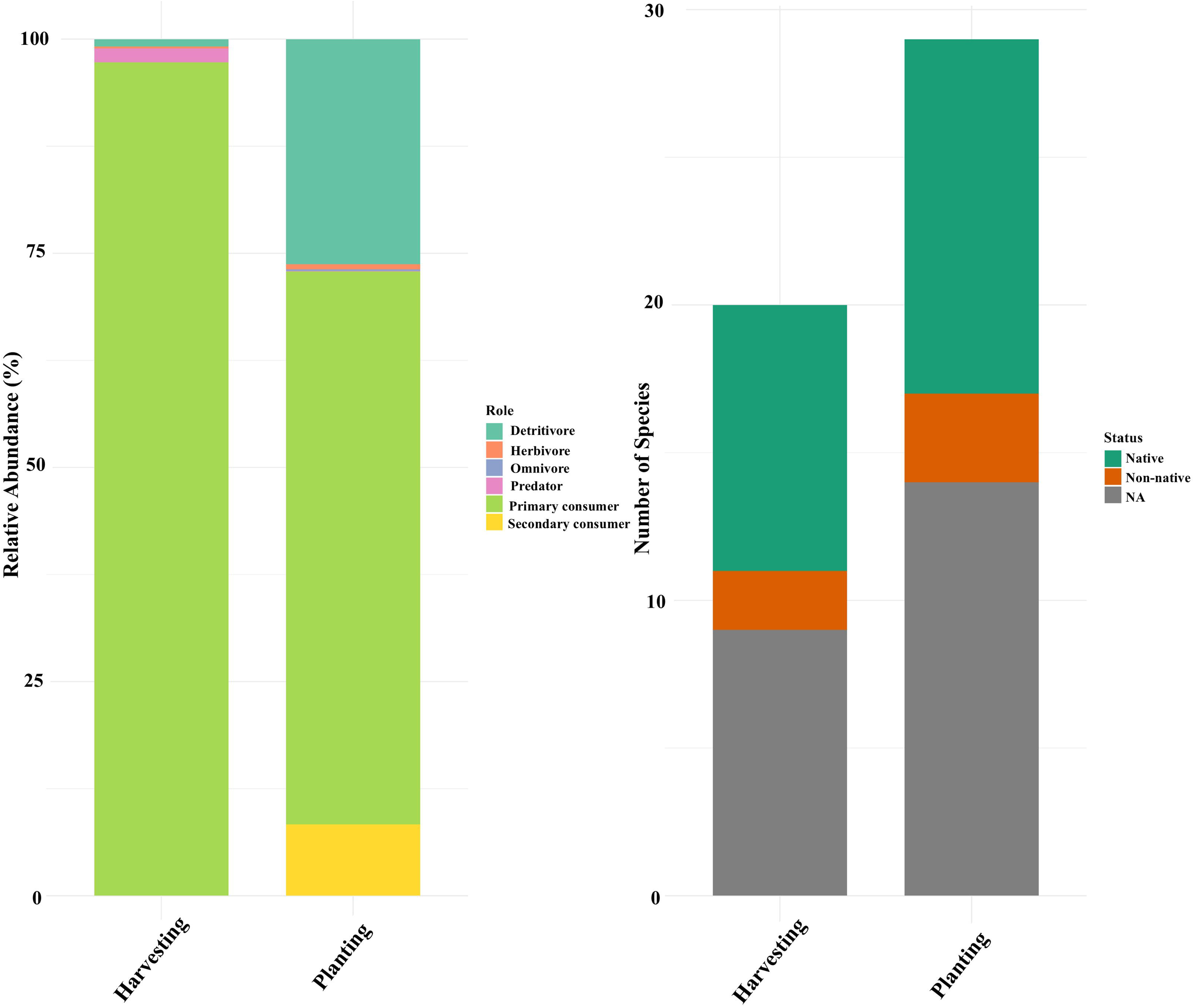
(A) Comparison of alpha diversity (observed ASVs) between field types (harvesting, orange; planting, green) across sites (n=8). (B) Non-metric multidimensional scaling (NMDS) plot based on Bray–Curtis dissimilarity showing beta diversity of communities between planting (green) and harvesting (orange) rice fields detected from eDNA metabarcoding. Each point represents a site, with ellipses indicating 95% confidence intervals for each field type.

**Figure 5.**
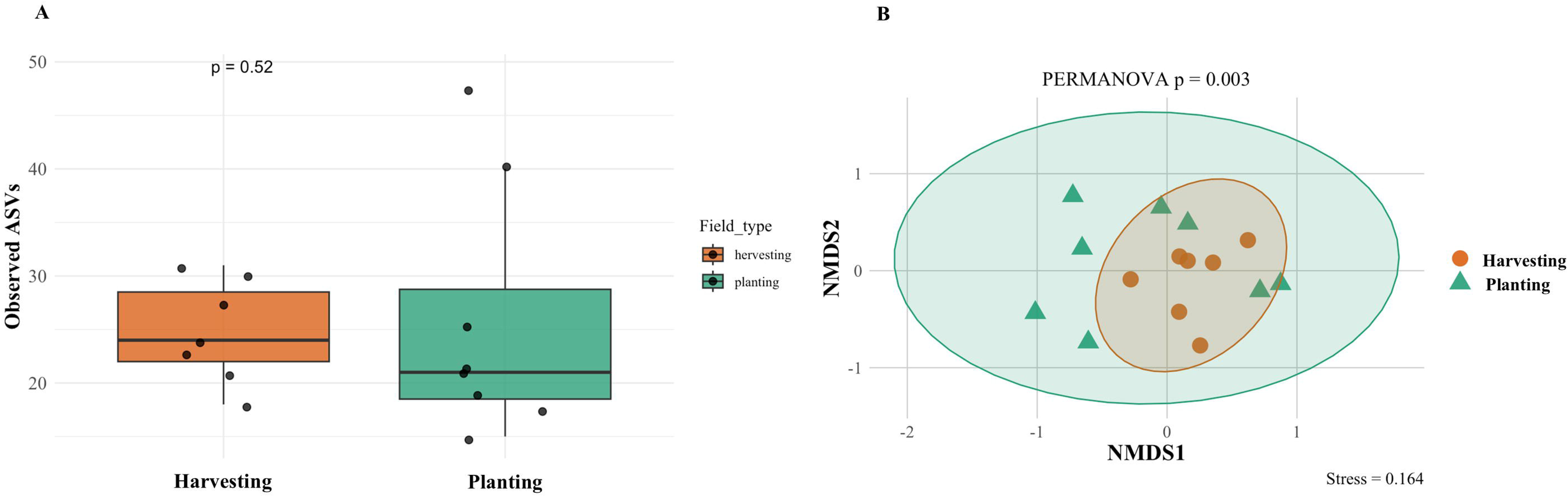
Functional roles and species status across rice field types. (Left) Relative abundance of species categorized by functional roles during harvesting and planting. (Right) Species richness grouped by published literature and the Catalog of Life in Taiwan (https://taicol.tw) (Table 4; native, non-native, and undermined/NA).

Despite no significant differences in alpha-diversity across cultivation times, the non-metric multidimensional scaling (NMDS) showed a pattern of differing community composition between cultivation times (Figure 5B) (stress = 0.164). The PERMANOVA test confirmed that the difference between the two growth phases was statistically significant (p = 0.003). Additionally, no significant differences were observed between native and non-native taxa across the two cultivation phases (Wilcoxon test, p > 0.99).

## Discussion

In this study, we aimed to establish eDNA metabarcoding as an effective method for detecting the vast array of interactive animal species from rice fields. Overall, the eDNA-based method successfully detected a wide range of taxonomic groups during both planting and harvesting phases, even with minimal sampling time and effort (Figure 3B, Table 4), consistent with the first hypothesis (H1). We successfully detected rice pests, including *Chilo suppressalis* (native), *Coptotermes formosanus* (native), and *Pomacea canaliculata* (non-native), among others (Table 4). Our results indicate that changes in rice cultivation phases correspond to significant changes in community composition (Figure 3A, B), though the species richness between the phases remains relatively static (Figure 5A). We didn’t observe any significant difference between native and non-native taxa across the two cultivation phases (Figure 4), counter to our second hypothesis (H2). However, this may also be a consequence of the relatively small sample size and limited study area. Future studies incorporating a larger number of samples and broader spatial coverage would improve the robustness of the analysis and may reveal patterns that were not detectable in the present study.

### Functional role of the detected organism

Our analysis of the rice field ecosystem revealed a diverse community spanning multiple trophic levels, including primary consumers, detritivores, predators, omnivores, and native and non-native agricultural pests (Table 4). Primary producers included rice crops and various phytoplankton. Detritivores such as *Metaphire posthuma* (Tsai et al., 2020), aquatic oligochaetes (*Allonais*, *Dero*, *Pristina*), soil invertebrates (e.g., *Homidia socia*, *Coptotermes formosanus*) (Cheng et al., 2022; Blumenfeld et al., 2021), and gastropods (*Melanoides tuberculata*) have an important role in nutrient cycling (Chang, 1991). Phytoplankton grazers included cladocerans (e.g., *Moina macrocopa*, *Ceriodaphnia cornuta*), rotifers (*Brachionus*, *Keratella*), and gastropods (*Pomacea canaliculata*, *Austropeplea ollula*) (Table 4). Omnivores such as ants (*Monomorium pharaonis*, *Pheidole* sp.) (Wetterer, 2010), and birds (*Passer montanus*, *Acridotheres tristis*) were also detected (Ding et al., 2020). In this context, birds may play a potential role as insect predators (Sottomayor et al., 2024). Secondary consumers included *Thermocyclops decipiens* and *T. taihokuensis* (Reid, 1989), while predators such as frogs (*Microhyla fissipes*, *Fejervarya* sp.) (Jang-Liaw & Chou, 2015) and aquatic insects (*Diplonychus esakii*, *Microvelia douglasi*) further structured the food web (Rieger, 1995; Ye et al., 2014).

Two taxa of particular concern, *Chilo suppressalis* and *Pomacea canaliculata*, are major rice pests responsible for economic damage (Chu, 1971); however, the detection of *Coptotermes formosanus* is also a concern, as they may damage the rice plants (Li et al., 2009). We also recorded high abundances of *Culex tritaeniorhynchus* (Figure 2), a vector of Japanese encephalitis virus, which may raise concerns about the management of stagnant water in rice fields (Table 4).

### Planting vs harvesting phase

The NMDS results demonstrate that arthropod community structure differs significantly between harvesting and planting phases. The tighter clustering of harvesting compared to the planting phase suggests more homogeneous community assemblages under harvesting conditions, possibly due to low availability of water or selective pressures associated with crop removal (Flannagan & Marshall, 2012; Saksongmuang et al., 2024). In contrast, the broader spread of identified ASVs during planting phases indicates greater community heterogeneity, which may reflect colonization dynamics, variability in early vegetation structure, or differences in resource availability (Mukherjee & Khan, 2017; Saksongmuang et al., 2024). This pattern is also supported by the measured physicochemical parameters, with the planting phase showing higher DO, ORP, conductivity, and TDS compared to the harvesting phase, possibly due to longer water residence time and more biological activity, whereas the harvesting stage was characterized by relatively drier conditions and reduced water availability (Table 6). However, in this study, the relationship between environmental parameters and indicator taxa could be better resolved with more robust seasonal sampling.

**Table 6.** The physicochemical properties and heavy metal concentration of water at agricultural fields during harvesting and planting. p-value are from Welch’s two-sample t-test (α=0.05). a and b subscripts represent significant differences.

| Parameter | Harvesting (Mean ± SE) | Planting (Mean ± SE) | p (Welch t-test) |
| --- | --- | --- | --- |
| pH | 7.52 ± 0.06 <sup>a</sup> | 7.54 ± 0.05 <sup>a</sup> | 0.7818 |
| mV pH | -21.48 ± 2.63 <sup>a</sup> | -26.70 ± 2.52 <sup>a</sup> | 0.2059 |
| ORP | 152.37 ± 11.67 <sup>b</sup> | 188.99 ± 3.77 <sup>a</sup> | **0.0270** |
| DO (mg/L) | 4.45 ± 0.02 <sup>b</sup> | 5.27 ± 0.26 <sup>a</sup> | **0.0197** |
| Conductivity (mS/cm) | 0.40 ± 0.04 <sup>b</sup> | 0.55 ± 0.05 <sup>a</sup> | **0.0363** |
| Resistivity (ohm·cm) | 3070.52 ± 382.50 <sup>a</sup> | 2111.05 ± 220.36 <sup>a</sup> | 0.0723 |
| TDS (mg/L) | 187.08 ± 20.39 <sup>b</sup> | 259.80 ± 22.12 <sup>a</sup> | **0.0424** |
| PSU | 0.26 ± 0.02 <sup>a</sup> | 0.25 ± 0.02 <sup>a</sup> | 0.7992 |
| Temperature (°C) | 29.32 ± 0.88 <sup>a</sup> | 26.08 ± 0.76 <sup>b</sup> | **0.0231** |
| Atm. Pressure (PSI) | 14.57 ± 0.10 <sup>a</sup> | 14.76 ± 0.02 <sup>a</sup> | 0.1412 |
| Fe (mg/L) | 0.15 ± 0.07 <sup>a</sup> | 0.03 ± 0.01 <sup>ab</sup> | 0.1676 |
| Li (mg/L) | 0.27 ± 0.00 <sup>a</sup> | 0.26 ± 0.00 <sup>b</sup> | **0.0082** |
| Mn (mg/L) | 0.31 ± 0.10 <sup>a</sup> | 0.24 ± 0.11 <sup>a</sup> | 0.6609 |
| Sr (mg/L) | 3.90 ± 0.87 <sup>a</sup> | 6.61 ± 1.09 <sup>a</sup> | 0.0938 |

The significant PERMANOVA results support our proposition that these biodiversity compositional differences are not random but instead driven by the phase of rice cultivation. This may highlight that agricultural practices, even across short time scales, can influence community composition, possibly through habitat modification, disturbance regimes, and resource distribution (Mukherjee & Khan, 2017). The harvesting phase encompasses warmer conditions compared to the planting phase, driven by warmer season and slight increases in soil pH and Pb, probably due to cumulative agrochemical deposition.

### Native and non-native status

Two frog species were identified as likely native to Taiwan. Among these, *Microhyla fissipes* is recognized as native and widely distributed (Jang-Liaw & Chou, 2015). The other taxon, *Fejervarya* sp., was identified only to the genus level, which includes both a native species (*Fejervarya limnocharis*) and a non-native species (*Fejervarya cancrivora*) present in Taiwan (Jang-Liaw & Chou, 2015; Lee et al., 2019). One native (*Passer montanus*) and an introduced bird (*Acridotheres tristis*) with widespread distribution in Taiwan was also detected from the rice field water (Ding et al., 2020). Among arthropods, *Coptotermes formosanus* is reported as a native pest species (Li et al., 2009; Blumenfeld et al., 2021); however, this species is considered among the top 100 invasive species in the world and its introduced population has been established in Japan, Hawaii, and the southeastern United States (Simberloff & Rejmanek, 2019; Blumenfeld et al., 2021). The native arthropod taxa detected included *Homidia socia*, *Culex tritaeniorhynchus*, *Sciara humeralis*, *Diplonychus esakii*, *Microvelia douglasi*, *Monomorium pharaonis*, and *Chilo suppressalis*. The non-native earthworm *Metaphire posthuma* was detected (Tsai et al., 2000); this species is now naturalized and does not appear to negatively impact the local environment. Additionally, one invasive snail (*Pomacea canaliculata*) which is responsible for various crop damage, and two native snails (*Austropeplea ollula* and *Melanoides tuberculata*), were detected together in the rice fields (Chang, 1991).

Although eDNA analysis of rice field water provided a comprehensive assessment of species presence, the array of biodiversity detection was still less than other previous conventional surveys conducted in Taiwan (Song & Kuo, 2022; Chen et al., 2023). In our results, we detected 77 animal ASVs, of which 34 were assigned to species level, which is lower than reported in previous conventional studies (Song & Kuo, 2022; Chen et al., 2023). The disparity with other studies may be due to lower sampling effort (16 pooled samples) in this study, where repetitive monitoring and an increase in sampling events can increase the detection (Yates et al., 2023). We did not explore the effects of eDNA persistence dynamics under hot climatic conditions and potential PCR inhibition (McCartin et al., 2022), but their influence cannot be neglected. Furthermore, universal concerns related to primer bias and limitations of reference databases require further investigation (Korbel et al., 2024). Taiwan has a good reference library (Hu et al., 2024), a direct comparison between the conventional detection and eDNA analysis can collect more evidence of disparity. Thus, future studies should implement a comparative analysis of conventional and eDNA methods to understand the potential community shift during cultivation and the robustness of eDNA biomonitoring compared to conventional approaches.

Fewer detections with eDNA sampling compared to conventional approaches may also suggest that the water sampling volume used here might not be sufficient to capture broad diversity (Cantera et al., 2019). Frequent clogging of filter papers from eDNA in turbid habitats, such as wetlands, is a key limitation (Hinlo et al., 2017). Future studies may look to test for pre-filtration (e.g., Turner et al. 2014), pooling of extractions from multiple smaller volume samples, or implement passive filtration methods (Stevens et al., 2024). Moreover, during the harvesting stage when less water is available, eDNA approaches may also create bias due to the increased inhibitors within the water samples, putatively impacting amplification and thus biodiversity recovery. Thus, additional soil and swabbing, or washing of rice plants for eDNA, could possibly maximize detection. Still, our eDNA-based approach detected a broad range of taxonomic groups with only limited sampling effort and without requiring extensive taxonomic expertise. Thus, our method provides a fast and effective tool for the regular monitoring of native and non-native pests in rice fields, and with developing technology, it will be easy to use for farmers. Furthermore, incorporating multiple sample types (e.g., water, soil, leaf swabs), using group-specific primers, and increasing the number of samples could further maximize the effectiveness of this monitoring approach.

## Conclusions

Environmental DNA metabarcoding is an effective tool in revealing the diversity and composition of communities in rice fields across Taiwan. Through this study, a wide range of taxa were detected in a limited number of sampling events, providing valuable insights into the pest community. Three detected pests (*Chilo suppressalis*, *Coptotermes formosanus*, and *Pomacea canaliculata*) are highly concerning. Notably, the presence of non-native *Pomacea canaliculata* dominates over native snails, raising concerns about the impact on native ecosystems and agricultural productivity. Furthermore, the detection of disease vectors like the Japanese Encephalitis virus from the stagnant rice field water is a rising concern for human health. These findings are particularly important for sensitive agricultural systems where ecological and biological processes are becoming increasingly recognized as essential for maintaining their ecological services. Overall, this study highlights the transformative potential of eDNA metabarcoding for rapidly evaluating dynamic biological and environmental systems.

## ACKNOWLEDGEMENTS

P.B. has been supported by Overseas Research Scholarships (ORS) from National Chung Cheng University as well as the Ministry of Education (MOE)- Industry- Academia project (Taiwan). The authors would also like to thank the Chiayi Christian Hospital (CYCH-CCU-2023-01) and National Science and Technology Council (NSTC:112-2116-M-194- 010) for financial support.

## DATA AVAILABILITY STATEMENT

Data sharing supporting the study is openly available in Dryad (Dataset http://datadryad.org/share/LINK_NOT_FOR_PUBLICATION/v8gjEC9xdWbgRxdx0GQ1BvlO bvD6UjM8EyZWVZOFdvg)

